# Connectome-mediated prediction of future tau-PET burden in Alzheimer’s disease

**DOI:** 10.1101/2020.08.11.246496

**Authors:** Pablo F. Damasceno, Renaud La Joie, Sergey Shcherbinin, Sudeepti Southekal, Vikas Kotari, Ixavier A. Higgins, Emily C. Collins, Gil D. Rabinovici, Mark A Mintun, Ashish Raj

## Abstract

Alzheimer’s Disease (AD) tau pathology originates in the brainstem and subsequently spreads to the entorhinal cortex, hippocampus and finally to temporal, parietal and prefrontal association cortices in a relatively stereotyped progression. Current evidence attributes this orderly progression to trans-neuronal spread of misfolded tau protein along the projection pathways of affected neurons. The aggregation of tau is being increasingly recognized as a trustworthy biomarker preceding the appearance of Alzheimer’s disease (AD) symptoms. One major goals of disease modifying therapies has been to stop or slow down the tau aggregation process. In order to evaluate drug efficacy, it would be desirable to have an accurate model predictive of a patient’s future tau burden, against which the tau measurements from drug-receiving cohorts could be compared. Here we report the development of such a model, evaluated in a cohort of 88 subjects clinically diagnosed as Mild Cognitively Impaired (MCI = 60) or Alzheimer’s disease (AD = 28) and tracked over a period of 18 months. Our approach combined data-driven and model-based methodologies, with the goal of predicting changes in tau within suitably specified target regions. We show that traditional statistical methods, allied to a network diffusion model for tau propagation in the brain, provide a remarkable prediction of the magnitude of incremental tau deposited in particular cortical areas of the brain over this period (MCI: R^2^ = 0.65±0.16; AD: R^2^ = 0.71±0.11) from baseline data. Our work has the potential to greatly strengthen the repertoire of analysis tools used in AD clinical trials, opening the door to future interventional trials with far fewer sample sizes than currently required.

## Introduction

Alzheimer’s disease (AD) is a double proteinopathy defined by the accumulation of extracellular β-amyloid (Aβ) plaques and neurofibrillary tau tangles in the brain. The search for prognostic biomarkers in AD is an area of intense research. Currently, signs of AD can be detected via fluid biomarkers (cerebrospinal fluid homogenates such as CSF tau and Abeta-42, and, very recently, plasma (Thijssen et al. 2020)), imaging (structural MRI for atrophy, FDG PET for glucose hypometabolism, amyloid and tau PET) and cognitive tests (see Ehrenberg et al. 2020 for a review). However, incomplete understanding of pathophysiology of progression, insidious onset, clinical heterogeneity and variability in speed and pattern of progression combine to preclude rigorous prognosis.

The mounting evidence associating tau accumulation and progression of Alzheimer’s disease (Giannakopoulos et al. 2003; La Joie et al. 2020; Mattsson-Carlgren et al. 2020) combined with the failure of anti-amyloid therapies to slow down cognitive decline in AD patients has made tau the new target for many therapies (Lo et al. 2014). For drug developers, the ability to predict the longitudinal progression of tau can enormously impact the design of and patient selection for clinical trials. For patients, knowledge of what the future holds can empower them to make informed choices regarding lifestyle and potential interventions.

As flortaucipir (FTP) PET, the current FDA approved tracer for *in vivo* measure of pathological human tau, becomes readily available, researchers have focused on the relationship between tau accumulation and downstream effects such as additional tau deposition, increase in brain atrophy, and decline of cognitive skills. Notably, it has been shown that the location of tau – but not amyloid – in the brain is a good proxy for future development of atrophy (La Joie et al. 2020). Forecasting the pattern and magnitude of tau accumulation, however, remain a challenge. While some studies (Jack et al. 2018; Pontecorvo et al. 2019) have shown a significant association between magnitude of SUVr increase and the pattern of baseline tau burden, others (Harrison et al. 2019; Sintini et al. 2019) have shown FTP increasing in frontal areas that do not coincide with regions with major FTP binding at baseline in AD patients or even that only CSF tau (but not tau PET) predicts tau PET accumulation in cognitively impaired individuals (Mattsson-Carlgren et al. 2020). Variety of these results illustrates a knowledge gap around the factors leading to tau accumulation in particular regions of the brain. Furthermore, the ability of these statistical regression analyses to predict change in tau in individuals is currently too low to warrant their use in clinical trials. This is an important limitation since the clinical utility of prognostic biomarkers rests squarely on their ability to predict future changes with respect to baseline.

### Combining data-driven and model-based approaches for predicting longitudinal tau changes

Our approach to predict future tau PET burden from baseline data was to consider additional factors that may drive tau accumulation, namely the stereotyped progression of tau, e.g. the Braak staging model (Braak and Braak 1996). In this model, neurofibrillary tau tangles are first found in entorhinal cortex and hippocampus (stages I-II), then spread into the amygdala and basolateral temporal lobe (stages III-IV), followed by iso-cortical association areas (stages V-VI). Mounting evidence suggests that tau spreads following vulnerable fiber pathways rather than spatial proximity (Englund, Brun, and Alling 1988; Kuczynski et al. 2010). It appears that tau follows a so-called “trans-neuronal spread” mechanism whereby proteins misfold, triggering misfolding of adjacent same-species proteins, and thereupon a cascade along neuronal pathways via transsynaptic or trans-neuronal spread follows (Palop and Mucke 2010, 201; Frost et al. 2009; Jucker and Walker 2013; Clavaguera et al. 2009; Jucker and Walker 2011). Since these processes involve network-based spread, several network models have been developed to use this information for disease tracking and progression forecasting (Raj, Kuceyeski, and Weiner 2012; Zhou et al. 2012; Iturria-Medina et al. 2014; Raj et al. 2015).

In this work we consider whether information about network connectivity between brain regions can be used to accurately predict tau accumulation over time. We incorporate the Network Diffusion Model (NDM, Raj et al. 2012, 2015) of network spread based on trans-neuronal transmission of misfolded tau, within a systematic data-driven statistical approach for predicting longitudinal progression of tau. We hypothesize that by combining the above models of tau propagation with data-driven approaches, we can vastly improve predictions of longitudinal changes in tau in individual subjects.

### Driving goal: predictive model as a computational endpoint for clinical trials

Our ultimate goal is to derive a computational biomarker of tau change that may be used as a monitoring and tracking tool in clinical trials. In the clinical trial setting, a key objective is to develop a measurable endpoint against which a prospective drug’s effect can be assessed. Our objective therefore is to find a set of features, based on baseline data, that is maximally predictive of the pattern and magnitude of future tau accumulation.

### Study design and the Network Diffusion Model

Subjects (N=88) clinically diagnosed as either MCI (N=60) or AD dementia (N=28) underwent MRI and FTP-PET imaging at baseline and at 18-month follow-up. Florbetapir-PET imaging was also acquired at baseline. Demographics are presented in Table 1.

**Table 1.** Demographics.

|  | **MCI**<br>**(n = 60)** | **AD**<br>**(n = 28)** |
| --- | --- | --- |
| Age | 72 ± 9<br>[50, 92] | 76 ± 8<br>[59, 95] |
| Females | 28<br>(47%) | 16<br>(57%) |
| APOE4 carriers | 6<br>(10%) | 4<br>(14%) |
| FBP positive<br>(visual read) | 28<br>(47%) | 18<br>(64%) |

AD: Alzheimer Disease, MCI: Mild Cognitive Impairment, APOE: Apolipoprotein, FBP: Florbetapir.
Data is shown as mean ± SD [minimum, maximum] for continuous variables or n (% available data) for categorical variables.

In order to predict tau-SUVr change over 18 months, we use the Network Diffusion Model (NDM). In this model, the longitudinal change in tau PET over each individual brain region (henceforth referred to “longitudinal FTP” or simply ∆FTP SUVr) is proportional to the baseline tau in that region modulated by a network connectivity factor. This connectivity factor (known as the network Laplacian 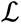) accounts for the diffusion of tau between connected region via fiber tracts (Fig. 1, top) and is a global parameter acquired from averaging diffusion tensor imaging (DTI) measurements from several healthy subjects (see Materials and Methods section for details). The proportionality constant (m) relating this modulated baseline tau and the longitudinal is a rate constant encapsulating the subject’s susceptibility to ongoing tau ramification. Since baseline tau and Laplacian are measurements available at baseline, the task of predicting longitudinal tau change becomes that of inferring m from baseline data.

**Figure 1.**
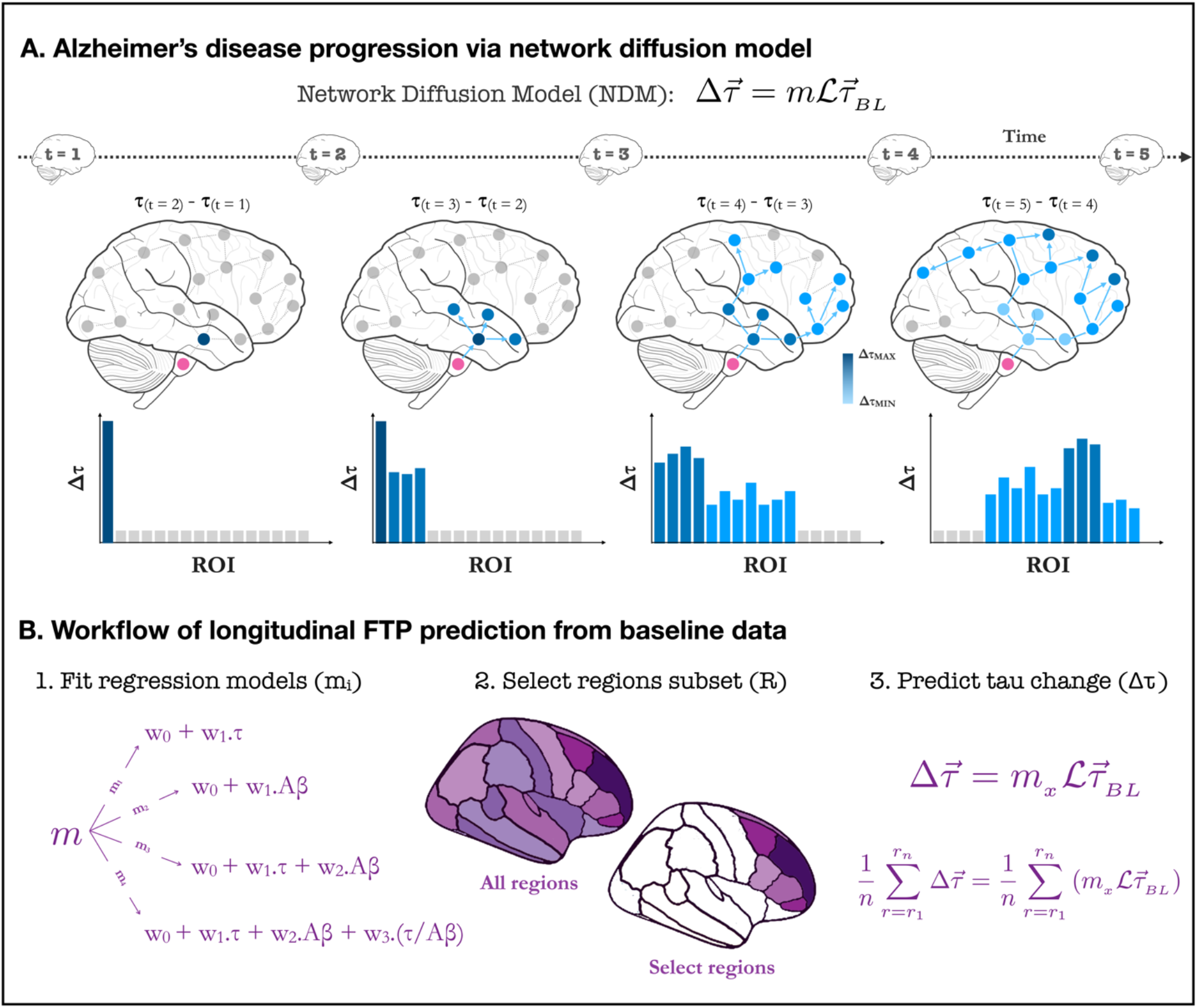
Longitudinal FTP prediction from baseline data via NDM. A. Schematics of tau progression following NDM. For each patient, the regional change in FTP between two scans (Δτ) can be modeled as the product of the patient’s susceptibility (m), the anatomically-generated graph Laplacian (L), and the regional pattern of deposition at first scan (τ). The timeline shows schematic tau patterns along the disease progression, followed by snapshots of tau change (Δτ) reflecting the difference between consecutive patterns. Starting in a seed region of interest (pink), tau tangles diffuse over time to nearby regions following anatomical connections. Darker areas indicate regions of higher change in tau over the measured time window. The histograms under each brain schematic show the evolution of Δτ over time in ROIs sorted by network proximity, highlighting how some regions previously showing elevated values of Δτ can become quiescent over time and vice-versa. B. Workflow for longitudinal FTP prediction. The first step for the regression is to choose a model relating baseline features and longitudinal tau change. Given a regression model, prediction can be made either at the whole-brain level (i.e. for every region of the brain) or for a select group of regions. NDM can also be used to propose such a select group. Once a target group of regions has been found, the model selected in step 1 can be applied to the NDM to generate a prediction of the longitudinal change in tau either region-wise on for the combined amount in a subset of regions.

Since predictions are made per region, it becomes trivial to rank these brain regions according to their predicted ΔFTP SUVr. A strategy we used in this work was to evaluate whether predictions in a subset composed only of highly burdened regions increased the prediction power of the model, compared to whole brain predictions.

Our workflow is the following (Fig. 1, bottom): we first test several regression models for m and measure their capacity to explain the variance in ΔFTP SUVr over 18 months in each individual brain region for each AD patient. We then compare this result against predictions of mean ΔFTP SUVr in *all* brain regions combined. Finally, we use NDM to identify subsets of ROIs expected to have high ΔFTP SUVr and test whether predictions in these “super ROIs” outperformed other empirically-derived “super ROIs” and whole brain predictions. For completion, all results were also compared against two other models: one identical to the NDM but ignoring the effects of network connectivity and another ignoring the effects of baseline data altogether.

To provide some insight into the role of m in our modeling, suppose we hypothesize a model m_A_ = w_0_.(age), meaning we assume that the relationship between modulated, regional baseline tau in region r for subject s 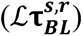 and the regional change in tau over 18-month period in this region is simply the patient’s age – in other words 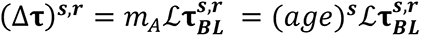. Since all terms in the right side are known and can be measured at baseline, we can determine the predictive power of our model by computing the explained variance between the model prediction 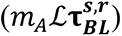 and the true measured change in tau ((Δ**τ**)^s,r^). Such explained variance is therefore an estimate on how well age predicts tau change over time.

In this work, we judiciously chose two covariates to construct m based on current models of neurodegenerative disease progression (Jack et al. 2010; 2013), that suggest that baseline tau and baseline amyloid should precede and influence future tau accumulation. Based on these two covariates, we proposed four simple regression models relating them to ΔFTP SUVr: m_1_ uses global baseline tau as the only covariate (Figure 2B, blue); m_2_ uses global baseline amyloid as only covariate (Figure 2B, brown); m_3_ uses a linear combination of the two prior covariates (Figure 2B, yellow), and m_4_ adds to this combination the ratio between the two covariates (Figure 2B, purple). The motivation for this last choice comes from recent findings that the colocalization of imaging biomarkers can boost signal-to-noise ratio and increase prediction accuracy in models classifying AD phenotypical variants (Damasceno et al. 2020).

**Figure 2.**
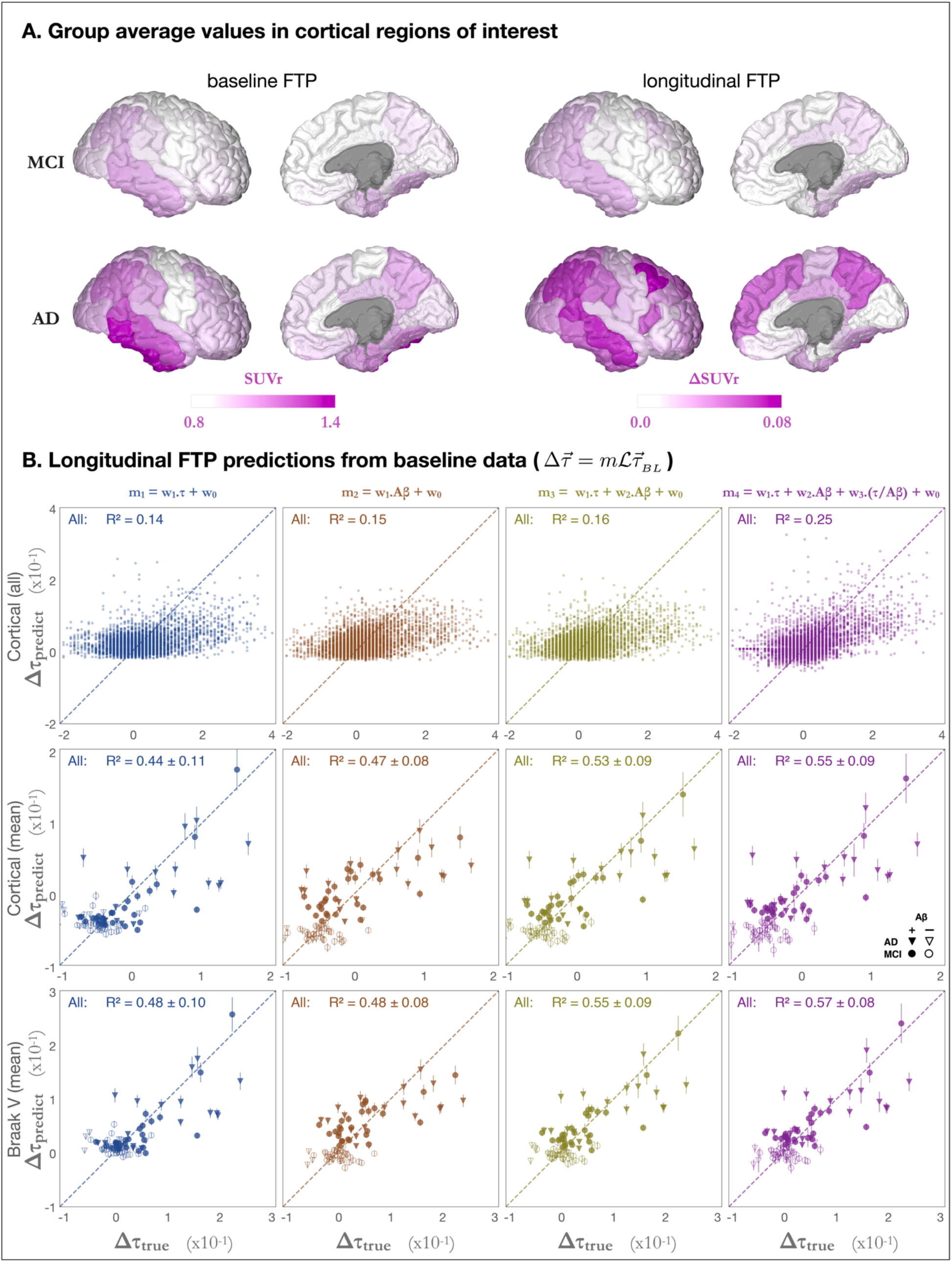
Longitudinal FTP predictions in selected ROIs. A. Group average baseline FTP-PET (left) and longitudinal FTP changes (right) for MCI (top) and AD (bottom) groups, shown in SUVr units. B. Longitudinal FTP predictions from baseline data. In each column, the longitudinal change in tau is modelled as a regression of different variables: based on baseline tau only (blue); baseline amyloid only (brown); baselines tau and amyloid (yellow); or baselines tau, amyloid, and the ration between them (purple). Top row shows the predictions for individual brain regions for all 28 AD patients (i.e. one dot per region per AD subject, total = 28 × 68), with a dashed line indicating the identity (i.e. prediction = true value). Second and third rows show mean ΔFTP SUVR over all cortical regions and all Braak V regions, respectively. Subjects with clinical diagnosis of AD (MCI) are shown as triangles (circles) whereas amyloid positive (negative) subjects are shown as solid (open) shapes. Mean explained variances and standard deviations were calculated over 5,000 bootstrap permutations.

## Results

### Measured ΔFTP SUVr is greater in regions with high baseline FTP in MCI individuals

Figure 2A shows FTP-PET data presented as parcellated (Desikan et al. 2006) tau-SUVr values normalized to white matter (see (Southekal et al. 2018) and Methods section for details). For the baseline data, both MCI and AD groups showed high FTP SUVr values in the temporal lobe, especially in the inferior temporal cortex. AD patients showed additional FTP signal in the parietal areas (both laterally and medially in the precuneus/posterior cingulate) and lateral frontal cortex. The pattern of longitudinal change in FTP strongly resembles that of baseline FTP in MCI, with most FTP increase being limited to the temporal and parietal lobe. In AD, higher FTP SUVr increase was observed in lateral temporal, parietal and frontal areas but some regions showing high tau at baseline (inferior temporal) do not show further tau accumulation, suggesting a “capping” behavior in agreement with previous findings (Schöll et al. 2016).

### Higher total cortical amyloid and total cortical tau at baseline lead to higher ΔFTP SUVr

We first predicted ΔFTP SUVr for every single individual brain region using the NDM predictor (see Eq 1 in Methods) (Figure 2B, top row), and achieved explained variances R^2^ = 0.14 and R^2^ = 0.15 for the models based on baseline FTP alone (m_1_) and Aβ alone (m_2_), respectively, whereas including both baseline predictors in a single regression (m_3_) marginally outperformed the model including Aβ only (R^2^ = 0.16). Including the ratio between FTP and Aβ as an additional feature (m_4_) significantly increased the explained variance (R^2^ = 0.25) compared to models m_1_, m_2_, or m_3_. For the m_3_ case, where we predicted tau change in individual brain regions, models of ΔFTP SUVr predictions that do not use NDM show slightly better or equivalent best explained variance compared to NDM models (See Figures SI 1-B and SI 1-C).

### Regression models predicting total longitudinal change in FTP over super-ROIs outperform predictions in individual regions

Inspired by the computer vision concept of super-pixels (Achanta et al. 2012), we defined super-ROIs as a collection of brain regions with similar features – here, similar predicted ΔFTP SUVr. Our hypothesis was that predictions over a collection of high signal ROIs could supersede those over individual, sometimes noisier regions. We initially defined two super-ROIs: the collection of all 68 Desikan-Killiani cortical regions, and the collection of 34 regions comprising Braak V stage. We then used the NDM to predict the mean longitudinal tau change, per subject, over each of these super-ROIs. Figure 2B (rows 2, 3) shows the values of the NDM prediction versus measurements of mean ΔFTP SUVr for each subject. Values were calculated by bootstrapping 5000 samples from the original data, from which mean and standard deviations for the explained variance were calculated. We note that our results were found to be robust against cohort selection criteria (see table S1). To facilitate this observation, data for the bottom two tests in Fig. 2B were displayed according to each subject’s amyloid statuses and clinical diagnose, although R^2^ values correspond to fitting *all* individuals at once, irrespective of their diagnosis or amyloid status. Similar to the individual region predictions case, the explained variances increase from model 1 to model 4 but are now much higher: R^2^ = 0.55 ± 0.09 for model 4 for cortical super-ROI and R^2^ = 0.57 ± 0.08 for Braak V super-ROI. These results suggested that applying the model over a collection of regions could increase the signal to noise ratio and boost prediction accuracy.

### Regression models predicting mean ΔFTP SUVr in NDM-derived super-ROIs outperform whole brain predictions

The previous result – that predictions over cortical and Braak V super-ROIs outperform those from individual regions – prompted the hypothesis that an even smaller subset of high signal regions could improve prediction accuracy. Selecting this subset of regions, however, is a non-trivial task: even splitting the 34 regions that make up Braak V staging into subgroups already leads to a factorial permutation explosion of impractical magnitude. To avoid this overfit-prone enumeration, we use a principled approach based on the NDM (Figure 3A). In short, our goal to reduce the number of regions in Braak V staging was achieved by first starting from an idealized version of an impaired brain seeded at Braak stages I-IV (i.e. τ = 1 for every region belonging to Braak stages I-IV as shown in Figure 3A-1); next we used the NDM to compute the longitudinal change in tau for all cortical regions (Figure 3A-2); we then ranked these regions according to such predictions and observed an elbow-like behavior, suggestive of a clustering of regions into a natural super-ROI (Figure 3A-3). Interestingly, the regions composing this super-ROI were found to be bilateral and were: Superior Temporal, Lateral Occipital, Precuneus, and Superior Frontal (purple in Figure 3A-3 and Figure 3A-4). As a final step in the workflow, we used the NDM to compute the mean longitudinal tau changes over the super-ROI and compared the results to the measured values. In what follows, we will focus on results for AD and MCI subjects. Results for all specific cohort subsets can be found in Table S1.

**Figure 3.**
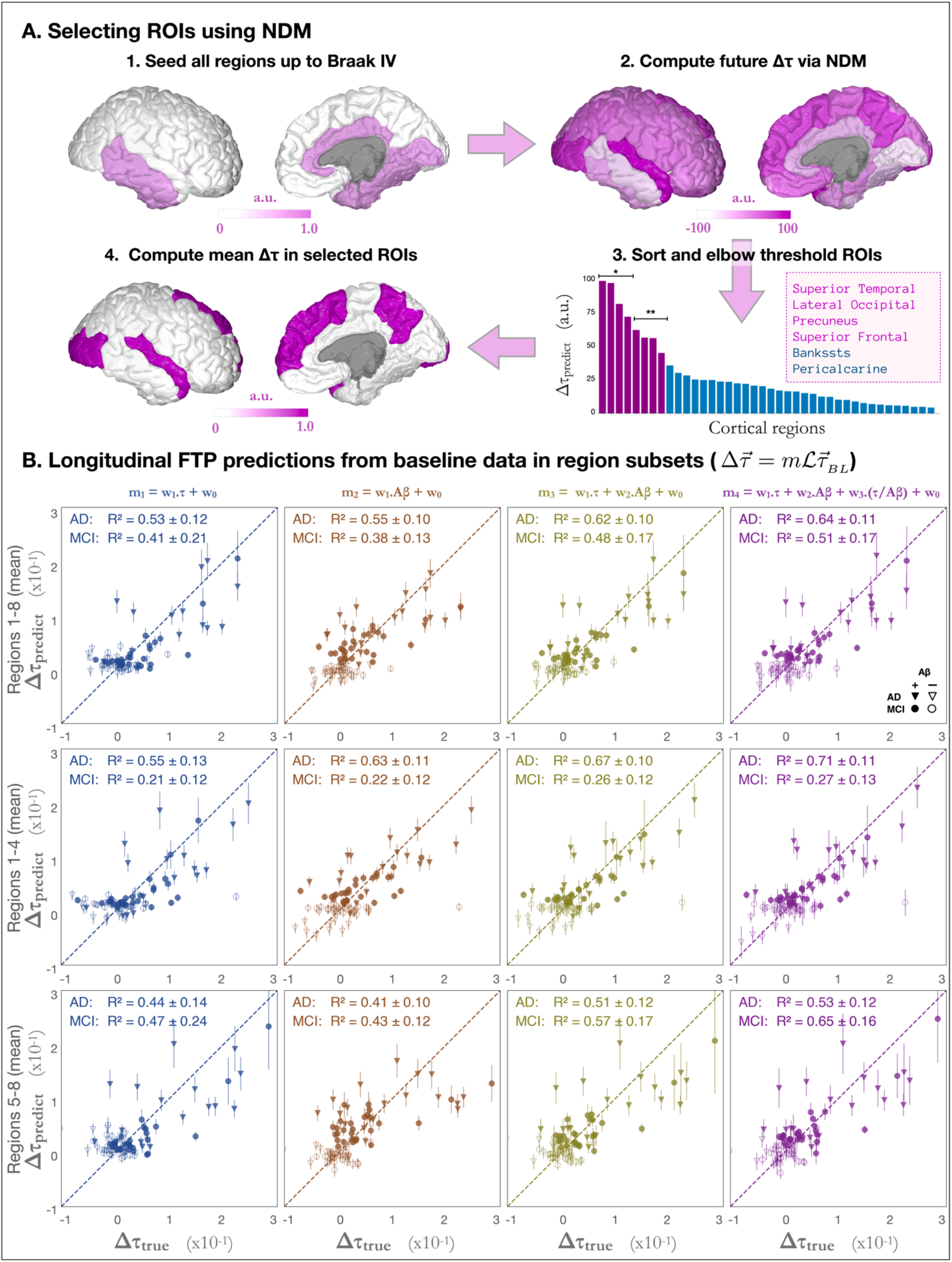
Longitudinal FTP predictions on NDM-derived brain regions. A. Workflow for regional subset selection using NDM. Starting with a brain “seeded” at Braak stage IV (highlighted regions in Figure 3A-1), NDM provides a snapshot with the pattern of future deposition (Figure 3A-2), allowing for a ranking of brain regions according to their predicted longitudinal burden (Figure 3A-3). After thresholding, a region subset containing top-8 regions (purple) is selected and the mean longitudinal tau change can be used as target for the regression predictions. * and ** in Figure 3A-3 represent subgroups used for predictions in cohorts with different clinical diagnosis. B. Prediction of future PET-tau from baseline values in NDM-derived. In each column, the longitudinal change in tau is modelled as a regression of different variables as in Figure 2B. Top, middle and bottom rows show predictions for NDM-selected regions 1-8 (purple in Fig. 3A-3), 1-4 (marked with * in Fig. 3A-3), and 5-8 (marked with ** in Fig. 3A-3) as explained in the text. Subjects with clinical diagnosis of AD (MCI) are shown as triangles (circles) whereas amyloid positive (negative) subjects are shown as solid (open) shapes. Mean explained variances and standard deviations were calculated over 5,000 bootstrap permutations.

Figure 3B shows that models predicting the mean ΔFTP SUVr across the super-ROI composed of the top eight regions were remarkably accurate (for AD: R^2^ = 0.53 ± 0.12, 0.55 ± 0.10, 0.62 ± 0.10, **0.64 ± 0.11** for models 1-4 respectively; for MCI: R^2^ = 0.41 ± 0.21, 0.38 ± 0.13, 0.48 ± 0.17, **0.51 ± 0.17**). As before, explained variance generally increased from models 1-4 with one exception for MCI patients, when model 2 outscored model 1.

While the predictions for AD across this super-ROI surpassed those obtained for all subjects across Braak V by over 10%, the results for MCI went in the opposite direction and decreased by 10% or more. This is to be expected since regions of high accumulation of tau in MCI subjects should not necessarily coincide with those in AD patients. To test whether subdivisions of this super-ROI could lead to cohort-specific prediction improvements, we further split the eight regions into two new super-ROI groups: regions 1-4, in the first, and regions 5-8 in the second.

Compared to results for the top eight regions, models trained to predict the mean ΔFTP SUr across the top four regions (Superior Temporal L/R; Lateral Occipital L/R – see Figure 3A-3*) showed a considerable increase in explained variance for AD patients (for AD: R^2^ = 0.55 ± 0.13, 0.63 ± 0.11, 0.67 ± 0.10, **0.71 ± 0.11** for models 1-4 respectively) accompanied by a drastic reduction in explained variance for MCI subjects (R^2^ = 0.21 ± 0.12, 0.22 ± 0.12, 0.26 ± 0.12, **0.27 ± 0.13** for models 1-4 respectively). Conversely, models trained to predict the mean ΔFTP SUVr across regions 5-8 (Precuneus L/R; Superior Frontal L/R – see Figure 3A-3*) showed the highest explained variance of all super-ROIs for MCI subjects, achieving R^2^ = 0.65 ± 0.16 for model 4 compared to R^2^ = 0.53 ± 0.12 for AD.

## Discussion

In this work, we developed a combination of data-driven statistical analysis and a model-based NDM approach to obtain a predictive tool for longitudinal tau change forecasting. To summarize our final result, our models achieve an explained variance of R^2^ = 0.71 ± 0.11 for AD subjects when predicting the mean ΔFTP SUVr over super-ROI composed of NDM-predicted regions 1-4 (L/R Superior Temporal and L/R Lateral Occipital), and R^2^ = 0.65 ± 0.16 for MCI subjects when using regions 5-8 (L/R Precuneus and L/R Superior Frontal). In both cases the selected regions come from the NDM model applied to a regional mapping corresponding to Braak stages I-IV. In both cases the prediction model incorporates the inter-regional tau spread process via the NDM applied to baseline regional tau. These results showcase how the NDM can be used not only as an accurate estimator of tau progression but also as an identifier of highly predictable ROIs. The cohort-dependency of our results is not only expected as it is also desired: ultimately, our model makes use of the different degrees of disease severity to identify the regions in which the rate of tau deposition will be severe and, at the same time, highly predictable.

### Comparison between NDM and non-Laplacian models

Figure S1 shows a comparison between three models relating regional baseline tau-SUVr and tau-SUVr change: the NDM, where tau-SUVr change is related to regional baseline tau-SUVr via a network diffusion term (A), one where this relationship is purely linear (B), and one where tau-SUVr change is independent of baseline tau-SUVr (C). Intuitively, these three models can be understood as local tau signal either: a) influencing nearby regions via network transport, b) causing local aggregation without transport, or c) having no relationship to future tau deposition. We found that the NDM model provides a similar prediction performance as the linear model, but that these models significantly outperform the independent model. Since the NDM model also incorporates local growth, this result implies that, in the short spam of 18 months, local aggregation is the most significant factor providing longitudinal accumulation of tau. The marginal improvements observed in some predictions, particular for super ROIs, suggest that connectivity-mediated diffusion will become more significant as data covering longer time periods become available.

### Relationship between amyloid and tau

AD is a dual proteinopathy. Although the topography of amyloid and tau depositions are distinct (Maass et al. 2019), there is a high correlation between the two pathological processes (Jack et al. 2019). It has been conjectured that for a given severity of amyloid beta, the severity of tau pathology can also be estimated (Brier et al. 2016). Our work goes one step further to provide a relationship between amyloid beta severity and magnitude and location of future tau PET burden. Our findings, showing a strong predictive association between total baseline amyloid and future tau PET accumulation support the scenario where amyloid is causally associated with the deposition of tau PET (Mattsson-Carlgren et al. 2020).

### Comparison with Machine Learning (ML) models

In this work we addressed two related but different problems: 1) how to predict the amount of tau aggregating over time in a subset of brain regions; and 2) how to optimally select such a subset of regions so to maximize our prediction capacity. ML has for quite some time been applied to many Alzheimer’s related questions, specially to the problem of predicting possible subject’s conversions to dementia (Davatzikos et al. 2011; 2009; Dickerson and Wolk 2013; Mattsson et al. 2014; Pachauri et al. 2011; Risacher et al. 2009). Deep Learning algorithms (Hinton 2007), which employ hierarchical learning via multi-level artificial neural networks, are particularly useful in this domain (Suk, Lee, and Shen 2014). While it is currently not known how well DL would perform on the problem being addressed here, we speculate based on a recent work applied to Parkinson’s disease (Nguyen et al. 2020) that DL will require far more samples in order to approach the accuracy we reported. Additionally, our usage of carefully constructed mix of prior information and biophysically-relevant models yield interpretability, which is missing in most “black box” DL approaches. Nevertheless, a rigorous comparison between these two approaches remains to be done and will form a key aspect of future work.

Similarly, recursive feature elimination and other machine learning techniques have been widely used for subset selection. These techniques, however, can lead to overfitting in small datasets like the one used here. The Network Diffusion Model provides a heuristic method to feature selection without the need for additional parameters, search loops, or data splits. Although the selected regions uncovered by the model are somewhat expected, the NDM offers not only a natural ordering of regions as well as a natural cut-off value for a small subset of regions expected to have high deposition of tau.

### Clinical and diagnostic implications

Data-driven methods are expected to revolutionize clinical development, from trial feasibility forecasting, to location and patient selection, to enrollment maximization (Shah et al. 2019). The ability to predict the longitudinal trajectory of a patient might allow for the creation of synthetic control groups (Abadie and Gardeazabal 2003; Abadie, Diamond, and Hainmueller 2010) for clinical trials. Our predictions rely on statistical regression models fitted to global burden of baseline tau and amyloid. Since these values are readily available at baseline, and given the high accuracy of our models, our results open the door for the use of predictive models of tau progression as an endogenous endpoint in clinical trials. Although the current focus was to develop endpoints to facilitate clinical trials, our work supports a key role of predictive models as an aid to the clinician, allowing them to predict what the patient’s neuroanatomic state will be in the future.

### Limitations and future work

This work has several limitations that should be acknowledged and addressed in future research. First, the observation window of 18 months is rather narrow in comparison to decades-long progression of the disease, precluding long-term longitudinal follow up. Second, because this kind of long-term tau-PET data are exceedingly hard to obtain, the current study used a very small sample size. A third possible weakness refers to the usage of group-level statistics. One formidable proposition for future work is that of identifying individualized regions where ΔFTP SUVr can be predicted with high accuracy, using subject’s baseline disease stage as opposed to their diagnose group. Another limitation related to SUVs is that SUVr itself can be vulnerable to biases, the biggest one being underestimation due to atrophy/partial volume effect. This could be a confound for model-based approaches that rely on SUVr. Fourth, this work used healthy reference rather than individual patients’ connectomes, as the available data did not include diffusion MRI protocols. It is likely that, compared to the quality provided by averaged reference connectomes individual patients’ connectomes are too noisy and confounded by image processing challenges to allow for meaningful analysis. Indeed, recent work suggests connectome variability exerts only minor influence on the NDM performance (Powell et al. 2018). It nevertheless remains to be seen whether individual connectomes can account for some of the unexplained variance in our data. Finally, one disadvantage of the NDM as a method for sorting brain regions is the necessity for a starting point (a seed) from which regions of highest ΔFTP SUVr are derived. As it became clear, one general model designed for AD patients (e.g. NDM seeded in Braak I-IV) does not work equally well for MCI subjects and vice-versa. One tantalizing implication of this necessity, however, is the possibility that, as data become available, cohort-dependent models – or even patient-tailored, bespoke predictions – become attainable.

## Materials and Methods

### Subjects

Baseline and 18-month follow-up flortaucipir and MRI images obtained in the A05 phase 2 study (NCT 02016560, (Pontecorvo et al. 2019)) were analyzed for N= 88 subjects who completed the study and were clinically diagnosed (Albert et al. 2011; McKhann et al. 2011) as either MCI (N=60) or AD dementia (N=28). Each subject’s T1-w MRI was processed using FreeSurfer V5.3. The same was repeated for the 18-month follow-up MRI. After the cross-sectional segmentation, the longitudinal pipeline within FreeSurfer was used to get better measurements of change. Each subject’s PET image was coregistered to the subject’s MRI in FreeSurfer space for every time point. The coregistered PET images are projected on to the FsAverage FreeSurfer surface and regional mean count values were extracted. Motion and acquisition start time-corrected PET images were aligned to the Freesurfer-conformed baseline MRI image for each subject. Regional mean count values were then extracted for 68 cortical regions and normalized to white matter (Southekal et al. 2018) to generate tau-SUVR values. Florbetapir images were also acquired and processed according to standard techniques (Joshi et al. 2015). Composite cortical SUVR values were used as covariates for 18-month tau-SUVR change predictions (see below).

### Network Diffusion Model (NDM)-Based Predictors of longitudinal Tau

The NDM provides a direct relationship between the (regional) values of baseline tau-SUVr and the (regional) change in tau-SUVr over time. Mathematically:

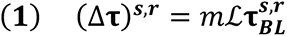

where (Δ**τ**)^s,r^ is the change in longitudinal tau for subject **s** in region **r**; **m** is the proportionality constant to be determined via regression (see following section below); **L** is the Laplacian of the structural connectome; and 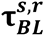 is the baseline tau for subject **s** in region **r**. In addition to predicting regional values, equation (1) above can also be used to calculate mean longitudinal values over a subset of regions as follows:

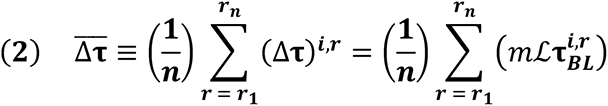

where the values in equation (1) are averaged over a subset of *n* pre-defined regions (e.g. all 68 cortical regions). Since both the Laplacian and baseline tau values are known, prediction of (Δτ)^s,r^ can be done via a simple regression for the value of **m**, as explained below.

### Regression Models

In order to predict (Δ**τ**)^s,r^ from equation (1) above, a proportionality constant **m** is needed. We tried several empirical models for m based on the available baseline data: model 1 uses total cortical tau, model 2 uses total cortical amyloid, model 3 uses a linear combination of these two features, and model 4 is the same as model 3 with the inclusion of a new feature, namely, the ratio between total cortical tau and total cortical amyloid (equations 3–6 below). The motivation for this last feature comes from recent findings that the colocalization of imaging biomarkers can sometimes boost signal-to-noise ratio and increase prediction accuracy (Damasceno et al. 2020).

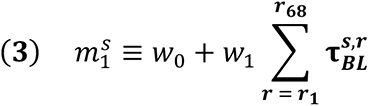

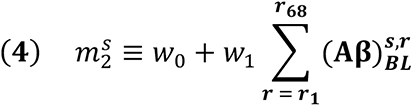

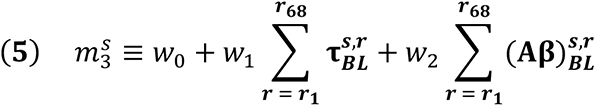

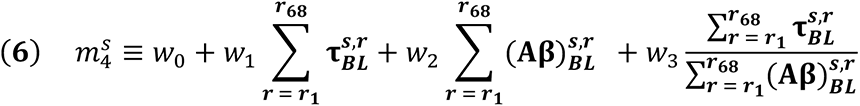

For each model above, we used Ordinary Least Squares (OLS) regression to find the best fit for the weight parameters **w**. When (Δ**τ**)^s,r^ was calculated as a mean over a subset of regions, bootstrapped was used to avoid overfitting (Fox and Fox 2016). In short, 5000 with-replacement permutations of subsets of the data were used. The mean and standard deviation values for the R^2^ reported in the figures were calculated over these 5,000 data subsets. When computing mean values over ROIs, values were normalized by regional area to allow for meaningful comparisons.

## Abbreviations

APOE: Apolipoprotein E
CSF: Cerebrospinal fluid
FTP: flortaucipir
PET: Positron emission tomography
MCI: mild cognitive impairment
PiB: Pittsburgh compound B
SUVr: standardized uptake value ratio

## Acknowledgements

We thank Dr. Ashley L. King for insightful discussions and Dr. Adam Fleisher for helpful comments.

## Study funding

AR, PFD were supported by NIH grants R01NS092802 and R01AG062196.

## Data availability

Data that support the findings of this study are available from the corresponding author upon reasonable request.

## Competing interests

**PFD, RLJ**, **AR** have no interests to declare. **SS, SS, VK, IH, EC, MM** are employees of Eli Lilly and Company. **GR** is Study Chair for the IDEAS study; Additional research support from National Institutes of Health (NIH), Alzheimer's Association, Tau Consortium, Avid Radiopharmaceuticals, and Eli Lilly. Consultant for Axon Neurosciences, General Electric (GE) Healthcare, Eisai, Merck. Associate Editor for JAMA Neurology.

